# Overmaturation as a major trajectory of naïve CD4 T cell aging

**DOI:** 10.64898/2026.02.02.703431

**Authors:** CJ Kelly, Janessa Wagandi, Danny Quaranta, Dina Prosser, Morgan Diegel, Nikolay Burnaevskiy

## Abstract

Aging is associated with progressive changes of cellular functional states, but whether these changes arise from novel programs or from distortion of normal developmental trajectories often remains unclear. Here, we investigate this question in the context of human immune aging by analyzing age-associated changes in CD4 T cells at single-cell resolution. We show that CD4 T cell subsets accumulate senescence-associated features at different rates and that the naïve CD4 T cell compartment is highly heterogeneous with respect to senescence markers. Strikingly, transcriptional changes associated with aging of naïve CD4 T cells parallel those observed during normal post-thymic maturation from recent thymic emigrants (RTE) to mature naïve cells. This similarity defines a transcriptional aging trajectory that we term overmaturation, characterized by exaggerated execution of a physiological maturation program. This overmaturation is consequential for T cell function, as we demonstrate through perturbation of transcription factor TOX, which is predominantly expressed in RTE and whose expression decreases with age. Together, our findings identify transcriptional overmaturation as a major trajectory of naïve CD4 T cell aging and suggest that aging-associated distortion of developmental programs can strongly contribute to immunosenescence.

## Main

With age, cells within tissues may change their functional state and these changes contribute to age-related pathologies. For instance, accumulation of cells that entered a state of senescence exacerbates many pathologies of old age^1^, while a shift of hematopoietic stem cells toward a myeloid-biased differentiation state can compromise the immune system function^2,3^. A better understanding of age-associated changes in cellular functional states may illuminate the mechanistic causes of aging and indicate novel targets for intervention.

T cells are critical mediators of adaptive immunity, and their functional decline with age contributes to immunosenescence. The T cell compartment comprises functionally distinct subsets—naive, memory, and regulatory cells—each with different roles in immune responses. Remarkably, T cells show signs of cellular aging early in life. Compared with T cells from cord blood, T cells of young adults (under 30 y.o.) already exhibit increased activity of senescence-associated β-galactosidase (SA-β-gal) ^4^. These SA-β-gal^Hi^ cells have higher expression of cyclin-dependent kinase inhibitors p16INK4a and p21CIP1, and diminished proliferative capacity, indicating their senescent-like state ^4^. These early changes suggest that T cell aging starts long before what would be considered an old age, making it a potential early driver of organismal aging.

Among T cell subsets, naive cells are of particular interest for understanding immune aging. The output of naïve T cells from the thymus declines precipitously after adolescence, due to a process known as thymic involution ^5,6^ As thymic functional capacity declines, homeostatic proliferation and half-life of existing naïve T cells increase, in part through elevated expression of IL-2 receptor (CD25), allowing the body to largely maintain the number of circulating naïve T cells ^7^. As a result, the maintenance of the naïve T cell pool increasingly depends on the longevity and homeostatic proliferation of existing naïve T cells ^8^. This compensatory mechanism effectively masks substantial underlying changes in cell age, proliferative history, and functional state. Determining how naïve CD4 T cells remaining in circulation change with age is therefore critical for understanding the origins of immunosenescence.

An important unresolved question is how naïve CD4 T cells age at the cellular level. One possibility is that aging primarily reflects changes in population structure, such as the loss of recent thymic emigrants (RTE) and the accumulation of mature naïve cells. A second possibility is that naïve T cells progressively acquire features of cellular senescence or other dysfunctional states that are distinct from normal developmental programs. A third, largely unexplored possibility is that aging represents an exaggeration or dysregulation of normal physiological processes. Discriminating between these models is essential for understanding whether immune aging is driven predominantly by damage accumulation, altered lineage composition, or distortion of normal differentiation trajectories.

Modern advances in single-cell transcriptomics give an opportunity to directly address these questions by resolving heterogeneity within the naïve CD4 T cell compartment from individual organism and tracing how subtle cellular states change with age. Here, we applied large-scale single-cell RNA sequencing to understand processes of senescence among heterogeneous CD4 T cells. We first found that memory and regulatory T cells exhibit elevated markers of senescence relative to naïve cells, and that the naïve compartment itself is highly heterogeneous with respect to both senescence-associated markers p16INK4a and p21CIP1. By separately analyzing recent thymic emigrants and mature naïve cells, we uncover a transcriptional aging trajectory that closely parallels normal post-thymic maturation. We term this process transcriptional overmaturation, characterized by progressive loss of RTE-associated gene expression and exaggerated acquisition of mature naïve transcriptional features. We detect this shift both in RTE and mature naive cells. Finally, we demonstrate that reduced expression of RTE-associated transcriptional regulators, such as TOX, has functional consequences for naïve CD4 T cell signaling. Together, our findings identify overmaturation as a major trajectory of naïve CD4 T cell aging and suggest new conceptual and mechanistic frameworks for understanding immunosenescence.

## Results

### Heterogeneity of senescence within CD4 T cell compartment

To understand how aging reshapes the functional states of CD4 T cells, we first asked whether senescence-associated heterogeneity could be resolved at single-cell resolution across major CD4 T cell subsets, and whether different subsets exhibit distinct aging trajectories. Addressing this question serves two purposes: it provides a framework for interpreting age-associated changes within naïve CD4 T cells, and it validates our experimental and analytical approach by comparing our findings with established features of T cell aging.

Hence, we first focused on the total CD4 T cell compartment to analyze markers of aging and senescence. We used CD4 T cells obtained from two middle-aged donors (45 y.o. male and 40 y.o. female) and stained live cells for SA-β-gal activity, using C_12_FDG ^9^. After the staining, cells were immediately sorted: approximately 20% of the brightest and 20% of the dimmest cells were separated into C_12_FDG^Hi^ and C_12_FDG^Lo^ groups. We refer to these groups as SA-β-gal^Hi^ and SA-β-gal^Lo^ respectively (Figure 1a). In addition, we collected cells that were stained and passed through the sorter but were not separated based on C_12_FDG staining. These non-sorted cells, that have undergone the same stressful treatment associated with C_12_FDG sorting, provide a reference to determine if sorted populations represent distinct subgroups within the total CD4 T compartment also work as a suitable control for cells that undergo the stress. After sorting, all samples were immediately fixed with formaldehyde and prepared for single-cell RNA-sequencing (scRNA-seq) using combinatorial cell barcoding ^10^. Sequencing data were analyzed using scanpy ^11^. Figure 1b shows UMAP projections of cells from one of the donors. Consistent with the sorting scheme, SA-β-gal^Hi^ cells exhibited higher expression of GLB1, the gene encoding beta-galactosidase (Figure 1b, top right panel), confirming the efficiency of the sorting.

**Figure 1.**
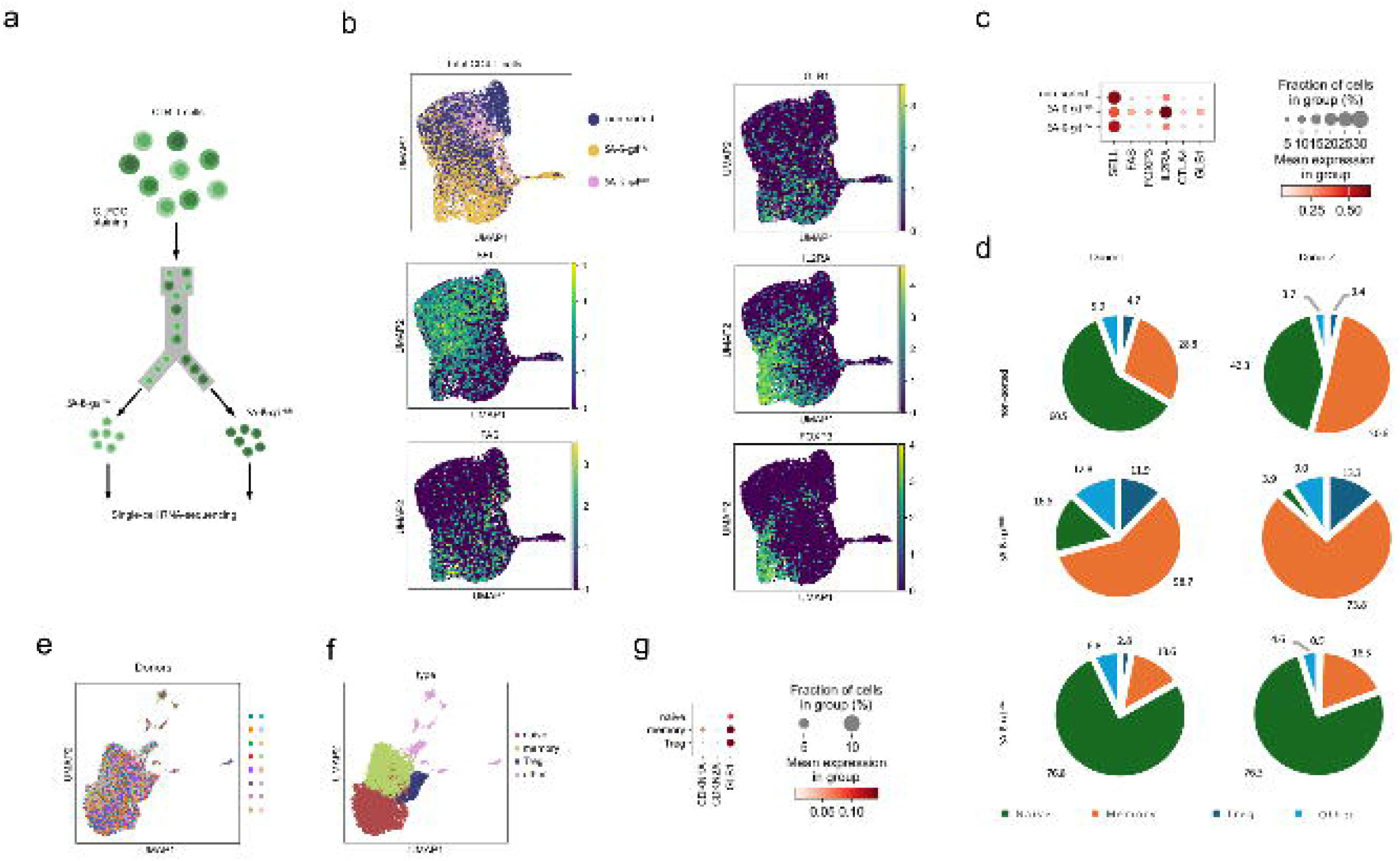
Senescence heterogeneity in CD4+ T cells. **a**. Experimental workflow: CD4+ T cells stained with C_12_FDG and sorted into SA-β-gal^Hi^ and SA-β-gal^Lo^ fractions for scRNA-seq. **b**. UMAP projections showing expression of GLB1, SELL, FAS, IL2RA, and FOXP3 across sorted CD4 samples. **c**. Expression of major CD4 cell types markers across sorted groups. **d**. Distribution of naïve, memory, and Treg cells in SA-β-gal^Hi^ and SA-β-gal^Lo^ samples. **e**. UMAP plots of scRNA-seq analysis of multiple donor samples. **f**. Cell type annotation of CD4 T cells from multiple donors. Annotation is based on markers shown in Figure S1c. **g**. Expression of senescence markers CDKN1A, CDKN2A, GLB1, in naïve, memory, and Treg populations from multiple donors.

Unsupervised embedding revealed that SA-β-gal^Hi^ and SA-β-gal^Lo^ cells occupied different regions within the total (non-sorted) population, suggesting that senescence-associated heterogeneity is structured rather than stochastic (Figure 1b, top left panel). We analyzed expression of the markers of naïve (SELL/CD62L), memory (FAS), and regulatory (IL2RA/CD25, FOXP3) T cells (see Figure 1b). Marker analysis indicated that SA-β-gal^Lo^ mostly represented naïve cells, while the SA-β-gal^Hi^ group mostly contained memory and regulatory T cells (see Figure 1b). The mean expression of the respective markers in different samples confirmed this conclusion (Figure 1c). We quantified the composition of each sample by counting cells that fell into respective Leiden clusters. This quantitative assessment confirmed that SA-β-gal^Hi^ is enriched with memory and regulatory cells, while SA-β-gal^Lo^ mostly contains naïve cells (Figure 1d). Similar results were obtained with the second donor (Figures 1d, S1a,b).

To determine whether these patterns generalize beyond the sorted samples, we analyzed scRNA-seq data from total CD4 T cells isolated from an independent cohort of 15 donors spanning a broad age range, without prior sorting by SA-β-gal activity. Figure 1e shows the resulting UMAP embedding. Cells were annotated as naïve, memory, or regulatory based on the same markers as before (Figure 1f, Figure S1c). Consistent with the earlier result, memory and regulatory cells had higher expression of GLB1 (Figure 1g). Furthermore, memory and regulatory T cells showed higher expression of CDKN1A (p21) and CDKN2A (p16), common markers of senescence (see Figure 1g). Hence, our initial conclusions made from the two C12FDG-sorted samples are generalizable, and we conclude that memory and regulatory T cells exhibit higher levels of classical markers of senescence and may follow different aging trajectories compared to naïve cells. These results are consistent with previous reports that Treg cells may senesce faster than conventional CD4 T cells ^12^. Higher SA-β-gal activity in CD4 memory cells compared to naïve cells is in agreement with recent findings that CD8 memory T cells exhibit higher levels of SA-β-gal than naïve CD8 T cells^4^.

First, these findings underscore the validity of our experimental approach for analyzing senescence of T cells. Results of our sorting approach and analysis are largely consistent with prior studies that analyzed SA-β-gal activity in T cells ^4,12^. Second, our data point to one of the factors that contributes to the increased abundance of SA-β-gal+ T cells with age, namely, the shift in the composition of the CD4 T cell compartment. With age, abundance of memory and regulatory T cells increases ^13,14^. Such a shift will subsequently lead to increased abundance of cells that exhibit signs of senescence, such as elevated SA-β-gal activity, and expression of p21 and p16.

Third, although SA-β-gal^Hi^ group was enriched with memory and regulatory cells, it also contained naïve cells, indicating that there is heterogeneity among naïve CD4 cells, with some falling into the SA-β-gal^Hi^ and others into the SA-β-gal^Lo^ group. After validating our approach for analyzing senescence heterogeneity, we wanted to determine next how SA-β-gal^Hi^ cells emerge among naïve CD4 T cells and how distinct naïve states relate to aging.

### Heterogeneity of senescence among naïve CD4 T cells

The presence of SA-β-gal^Hi^ cells within the naïve CD4 T cell population suggested that naïve cells are not uniform with respect to aging-associated features. We therefore next asked whether the naïve CD4 compartment contains distinct transcriptional states that differ in senescence-associated properties and could represent substrates for age-related change.

To address this question we isolated naïve CD4 T cells from a middle-aged donor (48 y.o. female) and applied the same C_12_FDG staining and sorting approach to separate SA-β-gal^Hi^ and SA-β-gal^Lo^ naïve CD4 T cells, followed by scRNA-seq. Unsupervised clustering revealed that SA-β-gal^Hi^ and SA-β-gal^Lo^ naïve cells occupied partially overlapping but distinct regions of transcriptional space, indicating structured heterogeneity within the naïve compartment (Figure 2a). Leiden clustering identified four major transcriptional clusters, which were unevenly distributed between SA-β-gal^Hi^ and SA-β-gal^Lo^ samples (see Figures 2a,b).

**Figure 2.**
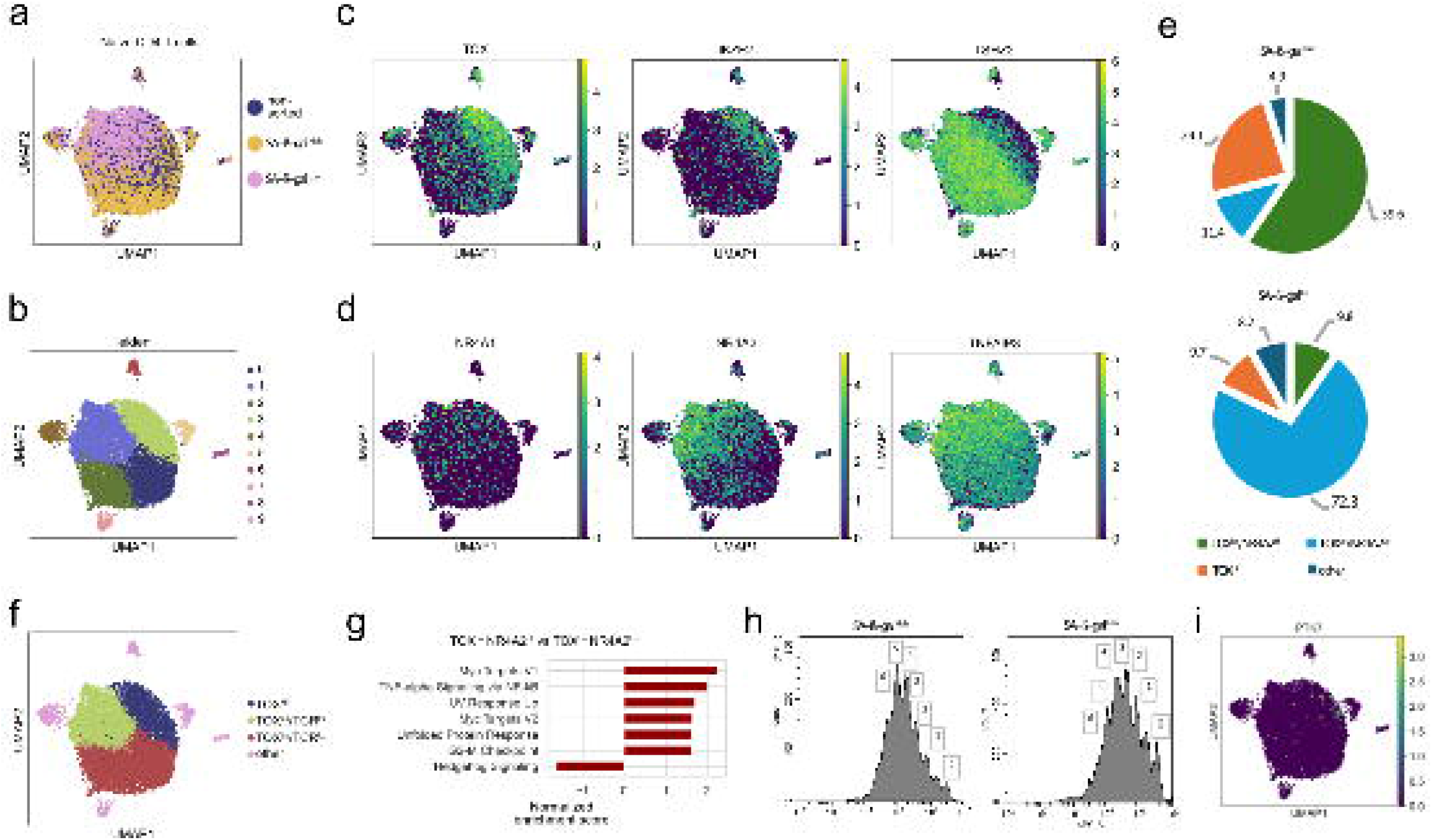
Heterogeneity of senescence within naïve CD4+ T cells. **a**. UMAP embedding of naïve CD4+ T cells sorted based on C_12_FDG staining. **b**. Leiden clustering identifies distinct subsets within naïve CD4 compartment. **c**,**d**. Marker expression identified two axes of cell heterogeneity. Cells can be separated as TOX^Hi^ vs. TOX^Lo^ and as NR4A2^Hi^ vs. NR4A2^Lo^. **e**. Abundance of the identified cell groups in SA-β-gal^Hi^ vs. SA-β-gal^Lo^ fractions. **f**. Annotation of the identified subgroups of naïve CD4 T cells. **g**. GSEA identified elevated NF-κB signaling in TOX^Lo^/NR4A2^Hi^ compared to TOX^Lo^/NR4A2^Lo^ cells. **h**. Proliferation assay comparing SA-β-gal^Hi^ and SA-β-gal^Lo^ naïve T cells. **i**. UMAP plot showing PTK7 expression marking RTEs.

Differential expression analysis revealed that naïve CD4 T cell heterogeneity could be largely described by two orthogonal transcriptional axes. One axis was defined by reciprocal expression of *TOX* and *TSHZ2*, separating TOX^Hi^ and TOX^Lo^ cells (Figure 2c). A second, independent axis was defined by expression of *NR4A1, NR4A2*, and *TNFAIP3*, separating NR4A2^Hi^ and NR4A2^Lo^ cells (Figure 2d). Importantly, these axes were orthogonal: both TOX^Hi^ and TOX^Lo^ populations contained NR4A2^Hi^ and NR4A2^Lo^ cells (see Figures 2c,d), indicating that naïve CD4 T cell heterogeneity reflects at least two separate biological dimensions.

The distribution of these states differed markedly between SA-β-gal fractions. TOX^Hi^ and TOX^Lo^/NR4A2^Hi^ cells were enriched in the SA-β-gal^Lo^ fraction, whereas TOX^Lo^/NR4A2^Lo^ cells were preferentially enriched among SA-β-gal^Hi naïve cells (Figure 2e). These results indicate that senescence-associated β-galactosidase activity within naïve CD4 T cells is not uniformly distributed but instead associates with specific transcriptional states. Similar patterns were observed in an independent donor, confirming the reproducibility of these findings (Figures S2a-e).

After identifying markers of the subgroups that exhibit different levels of SA-β-gal, we wanted to test if these subfractions are detectable without cell treatments associated with SA-β-gal staining and sorting. We revisited our scRNA-seq data from 15 donors and specifically focused on the naïve cell compartment (Figure S2g, also see Figures 1e,f). We found that separation on TOX^Hi^/TSHZ2^Lo^ versus TOX^Lo^/TSHZ2^Hi^ and NR4A2^Hi^ versus NR4A2^Lo^ can be detected even without sorting (Figure S2h). Hence, the identified fractions exist even in the unperturbed naïve CD4 compartment.

We then analyzed the distinguishable groups of naïve CD4 T cells. The NR4A2-defined axis suggested variation in basal T cell receptor (TCR) signaling, as *NR4A1* and *NR4A2* are immediate early genes induced by tonic TCR engagement (Figure 2f) ^15^. Consistent with this interpretation, gene set enrichment analysis showed elevated hallmark TNF-alpha signaling via NF-kB in TOX^Lo^/NR4A2^Hi^ cells compared with TOX^Lo^/NR4A2^Lo^ cells (Figures 2g, S2e). To test whether these transcriptional differences correspond to functional differences, we assessed proliferative responses following activation. We again sorted naïve cells into SA-β-gal^Hi^ and SA-β-gal^Lo^, then stained them with proliferation marker CFSE, and activated them *in vitro*. After 5 days, cell proliferation was examined with flow cytometry. Consistent with our hypothesis, SA-β-gal^Hi^ cells that are heavily enriched with NR4A2^Lo^ cells, proliferated better than SA-β-gal^Lo^ cells that are enriched with NR4A2^Hi^ cells (Figure 2h), consistent with prior observations that higher basal TCR signaling dampens proliferative capacity upon stimulation ^16^. Hence, we conclude that NR4A2^Hi^ cells exhibit higher inflammatory signaling before activation but show reduced proliferation upon stimulation, and hence NR4A2^Hi^ and NR4A2^Lo^ division reflects heterogeneity in basal TCR signaling strength (see Figure 2f).

Notably, our results diverged from a prior study of CD8 T cells, where high SA-β-gal activity correlated with slower proliferation ^4^. While better proliferation of SA-β-gal^Hi^ cells is surprising, it is consistent with our data that SA-β-gal^Hi^ cells are cells with low basal TCR signaling. It is not yet clear why TCR^Lo^ cells exhibit elevated activity of SA-β-gal. It was reported previously that TCR signaling negatively regulates basal metabolism, such that TCR^Hi^ cells are less metabolically active^17^. Due to the known importance of metabolism in aging and senescence, higher basal metabolic activity of TCR^Lo^ cells may lead to elevated SA-β-gal. In the future, it will be informative to examine the consequences of high and low tonic TCR signaling on the senescence of CD4 T cells.

In contrast, the TOX-defined axis suggested differences in developmental state rather than signaling activity. TOX^Hi^ cells expressed multiple transcription factors previously associated with thymic origin and early naïve T cell identity, including TOX2, SOX4, TCF4, IKZF2, BCL11A, and AUTS2. Indeed, these transcription factors are all part of the gene set distinguishing recent thymic emigrants (gene set “LEE_RECENT_THYMIC_EMIGRANT”, GSEA Molecular Signature Database). Furthermore, we found that TOX^Hi^ cells show elevated expression of protein tyrosin kinase 7 (PTK7), a known marker of recent thymic emigrants (RTE) ^18^. Hence, we determined that TOX^Hi^ represent recent thymic emigrants. A recent study also reported that within naïve compartment expression of TOX and SOX4 indeed marks RTE ^19^.

Thus, heterogeneity of the canonical senescence marker within the naïve CD4 T cell compartment reflects at least two separable dimensions: variation in basal TCR signaling strength and variation in developmental state. TOX-defined variation in developmental state is of particular interest: RTE express markers of stemness (SOX4, TCF4) and cytoprotective transcription factor IKZF2 ^20^, suggesting that RTE and mature naïve cells may age differently. We therefore asked whether age-related transcriptional changes in naïve CD4 T cells align with the developmental axis.

### Transcriptional overmaturation of naïve CD4 T cells with age

Strong separation of recent thymic emigrants (RTE) from mature naïve CD4 T cells upon SA-β-gal sorting raised the possibility that aging-associated transcriptional changes within the naïve CD4 compartment may share similarities with normal post-thymic maturation rather than reflect the emergence of a distinct senescent state. To test this hypothesis, we examined how RTE and mature naïve CD4 T cells change transcriptionally with age.

We isolated naïve CD4 T cells from cohorts of younger and older donors and performed single-cell RNA sequencing (Figure 3a). Upon completing scRNA-seq analysis, we identified fractions of RTE and mature cells within each sample (Figure 3b). As expected based on thymic involution, RTE were abundant in younger donors but declined markedly with age. Given that RTE exhibit lower SA-β-gal activity than mature naïve cells, this age-associated shift in naïve T cell composition is expected to contribute to increase of senescence-associated markers within the naïve compartment.

**Figure 3.**
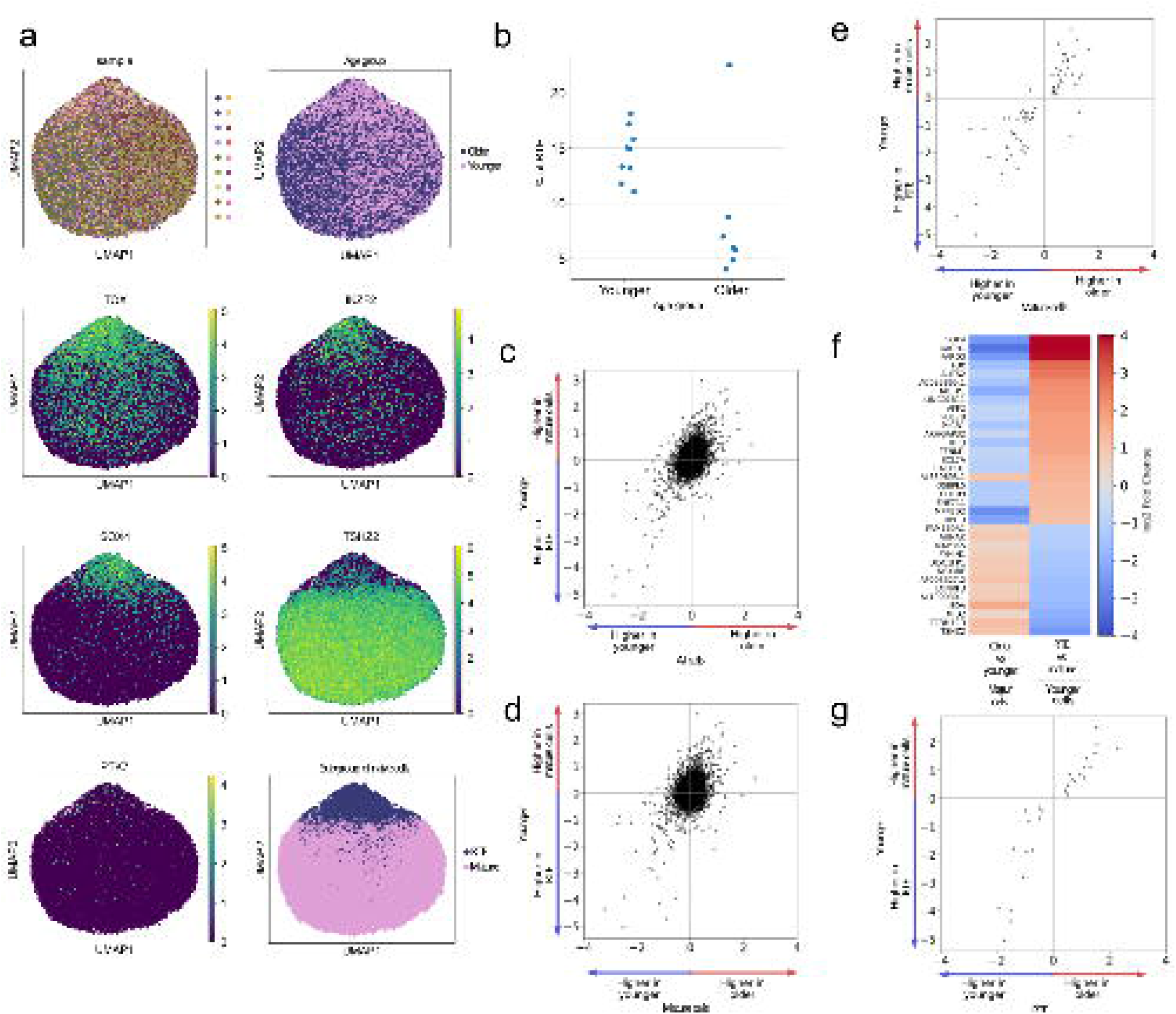
Transcriptional overmaturation of naïve CD4+ T cells with age. **a**. UMAP embeddings of naïve CD4+ T cells from younger and older donors. **b**. Quantification of RTE abundance with age. **c**. Transcriptional changes of total naïve CD4 T cells with age correlate with gene expression difference between RTE and mature cells. X axis shows relative expression of genes in younger and older cells as a log2 fold difference. Positive values correspond to higher expression of genes in older cells. Y axis shows relative expression of genes in RTE vs. mature naïve (only young donors included). Positive values correspond to higher expression of genes in RTE vs. mature cells. **d**. Transcriptional changes in RTE-to-mature transition and in mature naïve cells with age. X axis shows relative expression of genes in younger mature cells and older mature cells as log2 fold difference. Positive values correspond to higher expression of genes in older mature cells. Y axis shows relative expression of genes in RTE vs. mature naïve (only young donors included). Positive values correspond to higher expression of genes in RTE vs. mature cells **e**. Subset of genes from Figure 4d. Only genes that exhibited statistically significant change both during normal maturation in young donors and during aging of mature cells are included. **f**. Heatmap of genes showed in Figure 4e. Only genes with at least 2-fold difference between RTE and mature cells in young donors are included. Values correspond to log2 fold expression difference between younger and older mature cells, and log2 fold expression difference between RTE and mature cells from younger donors. **g**. Comparison of transcriptional aging changes during RTE-to-mature transition in younger donors and during aging of RTE. X axis shows relative expression of genes in young RTE vs. older RTE. Positive values correspond to higher expression of genes in older RTE. Y axis shows relative expression of genes in RTE vs. mature naïve (only young donors included). Positive values correspond to higher expression of genes in RTE vs. mature cells.

We first assessed transcriptional aging at the level of the total naïve CD4 T cell population. For each gene, we compared average expression between younger and older donors and asked whether these age-associated changes resembled transcriptional differences observed between RTE and mature naïve cells in young donors. Remarkably, these two sets of transcriptional changes were positively correlated (Figure 3c, Pearson correlation 0.38), indicating that genes downregulated with age tend to be enriched in RTE, whereas genes upregulated with age tend to be associated with mature naïve cells. Thus, at the population level, transcriptional aging of naïve CD4 T cells closely parallels the normal RTE-to-mature transition.

This similarity could, in principle, be explained solely by changes in population structure—namely, the loss of RTE with age. To determine whether mature naïve cells themselves undergo maturation-like transcriptional changes with age, we next restricted our analysis to mature naïve cells. Strikingly, age-associated transcriptional changes within mature cells again mirrored the transcriptional differences between RTE and mature naïve cells observed in young donors (Figure 3d). When focusing on genes that exhibited statistically significant changes during both during maturation or aging of mature cells, this relationship became even more pronounced (Figure 3e, Pearson correlation r -0.854, P-value 1.616e-21). Figure 3f shows the genes with the at least 2-fold difference between RTE and mature cells. Most of the genes associated with RTE in younger donors, decreased expression in mature cells with age, while expression of most of the genes associated with mature cells in young donors, increased further with in mature cells. These results indicate that mature naïve CD4 T cells continue to shift transcriptionally along the maturation axis with age, adopting an exaggerated mature gene expression program.

We considered the possibility that the observed transcriptional shift of mature cells is due to contamination by RTE. If our computational annotation of RTE and mature cells is not sufficiently precise, then the identified transcriptional changes associated with age can be simply caused by loss of RTE with age. However, we find that explanation not plausible. The fraction of RTE in the naïve compartment is insufficient to explain the observed magnitude of gene expression changes in mature cells. Even in young donors, RTE constitute about 20% of naïve cells (Figure 3b). Complete disappearance of RTE would increase the observed expression of genes associated with mature cells within the total naïve compartment by approximately 25%, which is not sufficient to explain the observed magnitude of transcriptional changes (see Figures 3e,f). Hence, in addition to overall reduction of RTE within the naïve compartment, mature cells exhibit signs of transcriptional overmaturation, with further loss of RTE markers and increased expression of mature cells markers.

We next asked whether RTE themselves exhibit similar aging-associated transcriptional shifts. Comparing younger and older RTE revealed that transcriptional aging of RTE is also highly correlated with RTE-to-mature transition (Figure 3g, Pearson r = -0.926, P-value = 6.630e-13). Thus, despite differences in developmental age and senescence-associated β-galactosidase activity, both RTE and mature naïve cells follow a similar trajectory of transcriptional aging.

Together, these findings indicate that transcriptional aging of naïve CD4 T cells comprises two complementary components: a reduction in the abundance of RTE and a progressive exaggeration of maturation-associated gene expression within both RTE and mature naïve cells. We term this process transcriptional overmaturation, reflecting a continuation of the normal post-thymic maturation program rather than the acquisition of a qualitatively distinct senescent state. These findings predict that genes with differential expression between RTE and mature cells may significantly affect CD4 T cell properties, including aging-relevant ones.

### Reduced expression of RTE-associated transcriptional program is consequential

It is reasonable to suggest that exaggeration of the mature naïve transcriptional state, and the concomitant loss of RTE-enriched regulators, should have functional consequences for naïve CD4 T cell behavior. The idea is indeed supported by prior studies. IKZF2 (HELIOS) is one of the genes that is enriched RTE compared to mature naïve cells. Decreased expression of IKZF2 (HELIOS) with age was shown to contribute to the increased inflammatory signaling of aged naïve T cells^20^. To test what other effects transcriptional overmaturation may have on the properties of naïve CD4 T cells, we focused on the transcription factor TOX. TOX is highly expressed in recent thymic emigrants but declines during post-thymic maturation and is further downregulated with age. TOX is a critical regulator of T cell development and function^21^. In addition, TOX regulates exhaustion of CD8 T cells, while in the CD4 lineage, TOX regulates T follicular helper development and modulates Th2 differentiation^22–25^. Altogether, prior literature pinpoint TOX as a versatile regulator of T cells development and functions^26^. Altered expression of TOX may therefore affect functional properties of naïve CD4 T cells upon activation.

We were specifically interested to test whether modulation of TOX expression in a physiologically relevant range would yield transcriptional changes consistent with known TOX biology. To this end, we employed epigenetic transcriptional modulators to activate TOX expression from endogenous locus. We synthesized epigenetic modifiers consisting of transcription activator-like effectors (TALE) fused to either KRAB domain or VP64 domain^27^. For additional controls, we used non-targeting TALEs, which should not bind any specific sequence within the genome, NT-KRAB and NT-VP64. All constructs carried a T2A-GFP tag to assess transduction efficiency and were delivered as lentiviral vectors.

We isolated naïve CD4 T cells from four healthy donors, activated them, and transduced with the designed TALE constructs (Figure 4a). GFP analysis indicated transduction efficiencies of 30–72% across samples. Upon harvesting, all samples were subjected to scRNA-seq. TALE-based activators increased average TOX expression in cells from all four donors compared to non-targeting constructs, while TALE-KRAB effectors reduced it, confirming that all effectors behaved as expected (Figure 4b). Importantly, expression of the related gene TOX2 was unaffected (Figure 4b).

**Figure 4.**
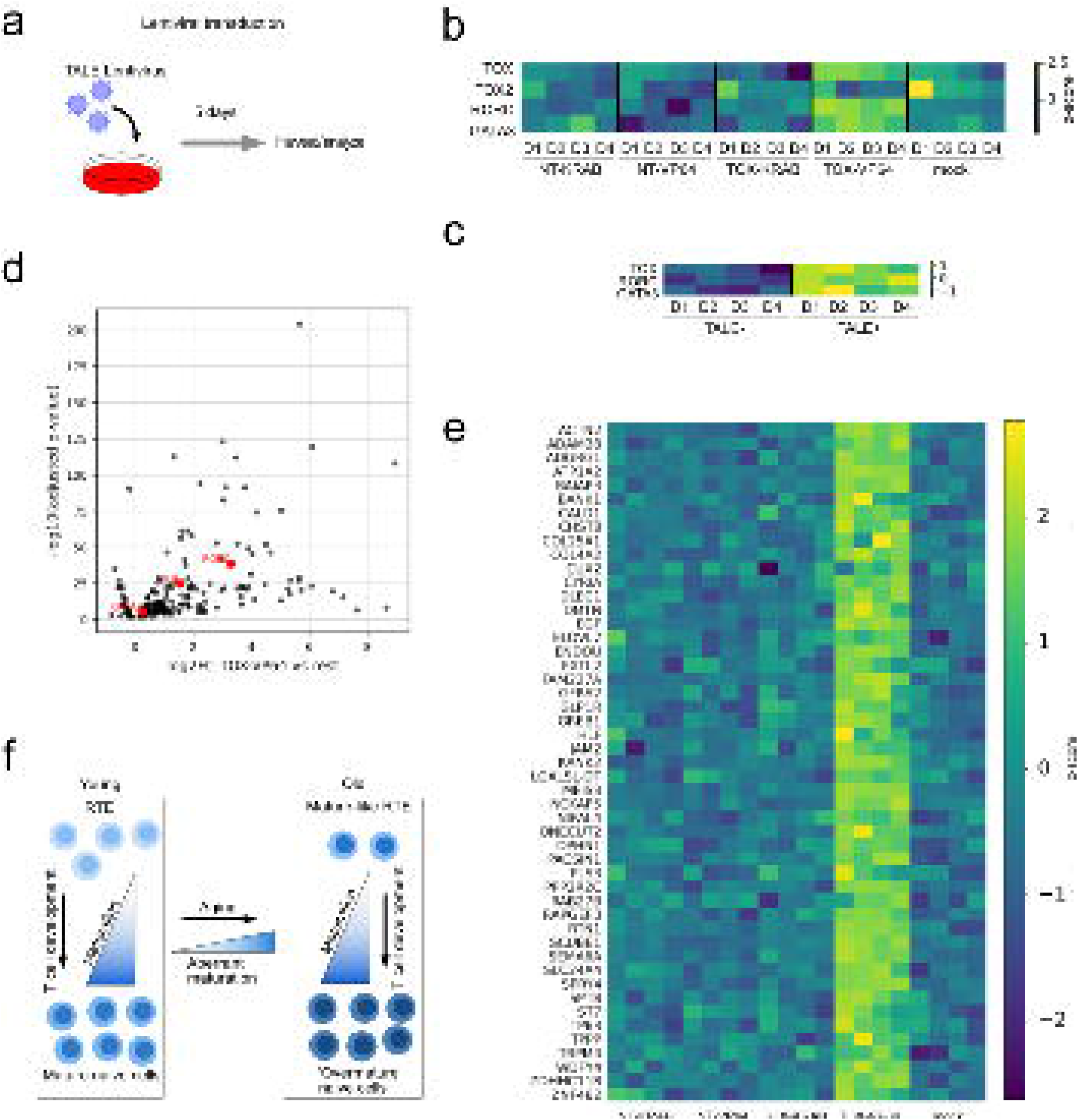
Activation of endogenous TOX upregulates lineage-determining transcription factors in naïve CD4 T cells. **a**. Experimental schematic. Naïve CD4 T cells were activated with anti-CD3/CD28, and transduced with lentiviral constructs encoding TALE-based epigenetic modulators. **b**. Expression of TOX, TOX2, RORC, and GATA3 across experimental conditions. TOX-VP64 increased TOX expression while TOX-KRAB reduced it relative to non-targeting controls. TOX2 expression was unaffected by any construct. RORC and GATA3 were specifically upregulated in TOX-VP64 samples. **c**. TOX, RORC, and GATA3 expression in cells with or without detectable TALE transcripts within TOX-VP64 samples. **d**. Genes with significant (adjusted p value <=0.05) expression change determined by three comparisons. **e**. Genes with significant expression change determined by three comparisons and at least 2-fold change in average expression across all comparisons. **f**. Model of overmaturation affecting naïve CD4 T cell properties.

We aimed to identify a robust transcriptional signature of TOX overexpression and avoid transcriptional noise and potential stress-response signals associated with cell handling and viral transduction. To this end, we performed three differential gene expression analyses: (1) TOX-VP64 samples versus all other samples; (2) TOX-VP64 versus TOX-KRAB samples; and (3) within TOX-VP64 samples, cells with detectable TALE transcripts versus those without. Cells with detectable TALE had significantly higher TOX expression (Figure 4c), indicating effective separation. Genes showing significant changes in all three analyses were retained (Figure 4d). We repeated identical analysis for TOX-KRAB samples, but it didn’t return any genes which showed consistent change in expression upon all comparisons. This lack of reliably affected genes in TOX-KRAB samples may result from incomplete repression of TOX and from relatively small fraction naïve cells that express TOX. We therefore focused on the genes affected TOX overexpression.

We examined the retained genes to determine whether TOX affects expression of differentiation regulators consistent with the known TOX biology. Previous studies indicated that TOX is critical for Tfh development, and modulates Th2 differentiation^25,28^. TOX also exerts a modest effect on Th17 differentiation^28^. Notably, even under non-polarizing conditions, TOX activation led to upregulation of GATA3 and RORC, critical regulators of Th2 and Th17 differentiation (Figures 4b). Induction of these factors was specific to cells with detectable TALE in TOX-VP64 samples (Figure 4c). Thus, upregulation of TOX expression from its endogenous locus is sufficient to alter transcription of fate-determining genes consistent with known TOX biology. We examined other genes that were strongly upregulated by TOX overexpression. The list includes genes involved in cytoskeletal and polarity remodeling (ACTN2, CALD1, PLS3, DMTN, SPTB, KANK2, OPHN1, PACSIN1, CYRIA, and TPPP) as well as membrane signaling molecules implicated in environmental sensing and cell positioning (ADGRG1/GPR56, GFRA2, GLP1R) consistent with TOX modulating early activation programs and potentially migratory behavior.

We conclude that differences in TOX expression between RTEs and mature cells, and between young and old naïve T cells in functionally relevant for CD4 T cell function. More broadly, these results support the idea that transcriptional overmaturation is consequential and may contribute to functional changes of the aging immune system. Our findings warrant further investigation into the role other RTE- and mature cells-associated transcriptional regulators in age-associated changes of naïve CD4 T cells.

## Discussion

Aging is accompanied by progressive changes in cellular functional states, yet the nature of these changes— and whether they reflect novel dysfunctional programs or distortions of normal differentiation trajectories— remains poorly understood for many cell types. Here, by resolving senescence-associated heterogeneity within the human CD4 T cell compartment at single-cell resolution, we identify transcriptional overmaturation as a major trajectory of naïve CD4 T cell aging. Rather than acquiring a qualitatively distinct senescent state, aging naïve T cells progressively exaggerate a normal post-thymic maturation program, characterized by loss of recent thymic emigrant (RTE)-associated transcriptional regulators and reinforcement of mature naïve gene expression.

A central implication of the overmaturation model is that age-associated loss of RTE-enriched transcriptional regulators may have functional consequences. We tested this prediction by focusing on TOX, a transcription factor highly expressed in RTEs that declines with both maturation and aging. Previous studies have established that TOX and the related factor TOX2 promote T follicular helper (Tfh) cell differentiation through a feed-forward loop with BCL6 and by remodeling chromatin accessibility at Tfh-associated loci^29^. TOX has also been implicated in modulating Th2 and Th17 differentiation^26,28^. We therefore asked whether restoring TOX expression could affect the differentiation potential of naïve CD4 T cells. Activation of TOX from its endogenous locus in primary human naïve CD4 T cells led to upregulation of differentiation-determining transcription factors, including GATA3, RORC, and IKZF3. These changes occurred under non-polarizing conditions, indicating that TOX primes cells for multiple alternative fates rather than driving commitment to any single lineage. Further studies should test how altered TOX expression affects functional properties of aged naïve T cells.

In addition, reduced expression of IKZF2 and subsequent alteration of STAT5 signaling is another documented feature of naïve CD4 T cell aging^20^. We found that IKZF2 is predominantly expressed in RTEs; hence, reduced presence of RTEs in circulation and reduced activity of the RTE-associated transcriptional program may both contribute to the observed decline of IKZF2 expression with age^20^. We expect that our data will help reconcile multiple previously described features of naïve CD4 T cell aging within a unified framework of transcriptional overmaturation.

Several important questions remain. The upstream drivers of overmaturation are not yet clear and may include increased cellular lifespan, cumulative proliferative history, or changes in systemic inflammatory and metabolic cues with age. It is also unknown whether overmaturation is reversible, or whether interventions that restore the RTE-associated transcriptional program can meaningfully improve immune function in older individuals.

In summary, our study identifies transcriptional overmaturation as a major trajectory of naïve CD4 T cell aging and demonstrates that age-associated loss of RTE-enriched transcriptional regulators may have functional consequences for T cell. These findings frame aging of naïve CD4 T cells as exaggerated execution of a physiological post-thymic maturation program rather than acquisition of a distinct senescent state and raise the possibility that similar exaggeration of normal developmental trajectories may underlie aging in other cell types.

## Methods

### Donor samples and T cell isolation

Peripheral blood mononuclear cells (PBMCs) were isolated from leukopacks provided by StemCell Technologies. All blood samples were collected from superficially healthy donors. Naïve and total CD4+ T cells were isolated using EasySep™ Human Naïve CD4+ T Cell Isolation Kit II or EasySep™ Human CD4+ T Cell Isolation Kit per manufacturer instructions (Stemcell Technologies). For age-comparison studies, younger/older matched pairs of cryopreserved PBMC were thawed and prepared in parallel for scRNA-seq; pairs were matched by gender and ethnicity, and BMI. Age range for younger donors: 20-26 years old, older donors: 48-60 years old. For scRNA-seq profiling of total CD4 T cells from 15 donors (Figure 1e), we used cells obtained from donors in the age range of 22-61 years old with median age of 33 years.

### SA-β-gal activity (C_12_FDG) staining and FACS

Senescence-associated β-galactosidase activity was measured with C_12_FDG. For the staining, cells were first pre-treated with 100 nM Bafilomycin A1 to alkalinize lysosomes following the manufacturer protocol. After pre-treatment, cells were incubated with 33 µM C_12_FDG (Thermo Fisher), then washed, stained with viability dye, and sorted by Astrios cell sorter. We collected: (i) SA-β-gal^Hi^ (top ∼15-20% of C_12_FDG fluorescence), (ii) SA-β-gal^Lo^ (bottom ∼15-20%), and (iii) non-sorted (C_12_FDG-stained, passed through the sorter without gating).

### Single-cell RNA sequencing

Cells were fixed with 1% formaldehyde, permeabilized, and subjected to combinatorial barcoding-based for scRNA-seq^10^. Upon completion of the barcoding procedure, short-read libraries were constructed as previously described ^10^ and sequenced on Illumina NovaSeq 6000 and NovaSeq X instruments. Downstream analysis was done in scanpy package ^11^. Unless noted, cells with very low complexity (determined by kneeplot) or high mitochondrial RNA (more than 10%) were excluded; genes detected in <10 cells were dropped. Counts were library-size normalized and log-transformed. Dimensionality reduction was done through PCA, followed by neighborhood graph construction, and UMAP embedding. Leiden clustering was used for unsupervised partitioning.

### Differential expression and statistics

Differential expression analysis of sorted cells was done using scanpy’s Wilcoxon rank-sum test with Benjamini-Hochberg correction. Effect sizes are reported as log fold-change on normalized counts. For sample composition plots, fractions were computed based on Leiden cluster annotations and expression of marker genes, as indicated in figure legends.

We performed differential gene expression analyses using a pseudobulk approach with DESeq2 (implemented via PyDESeq2). Raw UMI counts—without normalization or log transformation—were aggregated by summing counts across cells within the same biological grouping (e.g., sample, treatment, or TOX status). This generated one pseudobulk profile per group. For paired analyses, we retained only samples with matched conditions across comparisons. Statistical significance was assessed using Wald tests, and we adjusted p-values for multiple testing using the Benjamini–Hochberg procedure.

To reduce instability in fold-change estimates, we excluded genes with low average expression before downstream comparisons. All log2 fold changes are reported relative to the specified reference condition for each contrast.

### Gene Set Enrichment Analysis (GSEA)

For GSEA, its python implementation GSEApy was used. Genes were ranked using scanpy tools and Wilcoxon statistic test as described above. Rank lists of genes were used as inputs for GSEApy. For plotting GSEA results, only gene sets with adjusted *p* value of 0.05 or less were included.

### Cell proliferation assay

Naïve CD4+ T cells were sorted into SA-β-gal^Hi^ and SA-β-gal^Lo^ groups as described above, stained with CellTrace Far Red (Thermo Fisher), and cultured with or without CD3/CD28 Dynabeads stimulation following manufacturer recommendations (ThermoFisher). Proliferation was assessed after 5 days by flow cytometry, quantifying dye dilution across cell divisions. Population divisions which we defined by individual peaks of signal.

### TALE-based transcriptional modulation

To manipulate expression of TOX and TOX2, we engineered Transcription Activator-Like Effector (TALE) fused to either VP64 (activator) or KRAB-DNMT3A (repressor) domains. TALEs were designed to target the vicinity of target genes promoters, -500 to +150 bp relative to transcription start sites. We used TALE-based effectors in pools, 10 different effectors for each TALE RNA transfection, and 3 different effectors for TALE lentiviral deliveries. Within the pools, constructs differed from each other in the binding positions within the targeted promoters. For controls we used constructs with TALE sequences not specific to any region in the human genome. TALE effectors were delivered either as RNA or as lentiviral constructs. For transfection of TALE effectors RNA, naïve CD4 T cells from two donors were activated with CD3/CD28 Dynabeads overnight, and then electroporated using BTX system. We used 250k cells and 5ug of RNA per transfection. Samples transfected with RNA-based activators were harvested 2 days after transfection, and samples transfected with RNA-based repressors were harvested 4 days post-transfection. For lentiviral transduction, naïve cells from four donors were activated overnight with CD3/CD28 Dynabeads,and transduced at MOI ∼1. After overnight incubation, the transduction media was removed and cells were cultured for 5 days before harvesting.

## Data and code availability

Raw sequencing data, processed count matrices, and metadata (including donor age/sex, and run parameters) will be deposited to GEO. Analysis notebooks will be released at GitHub upon publication. TALE target sequences will be provided as accompanying materials for the paper.

## Figure Legends

**Supplementary Figure S1.**
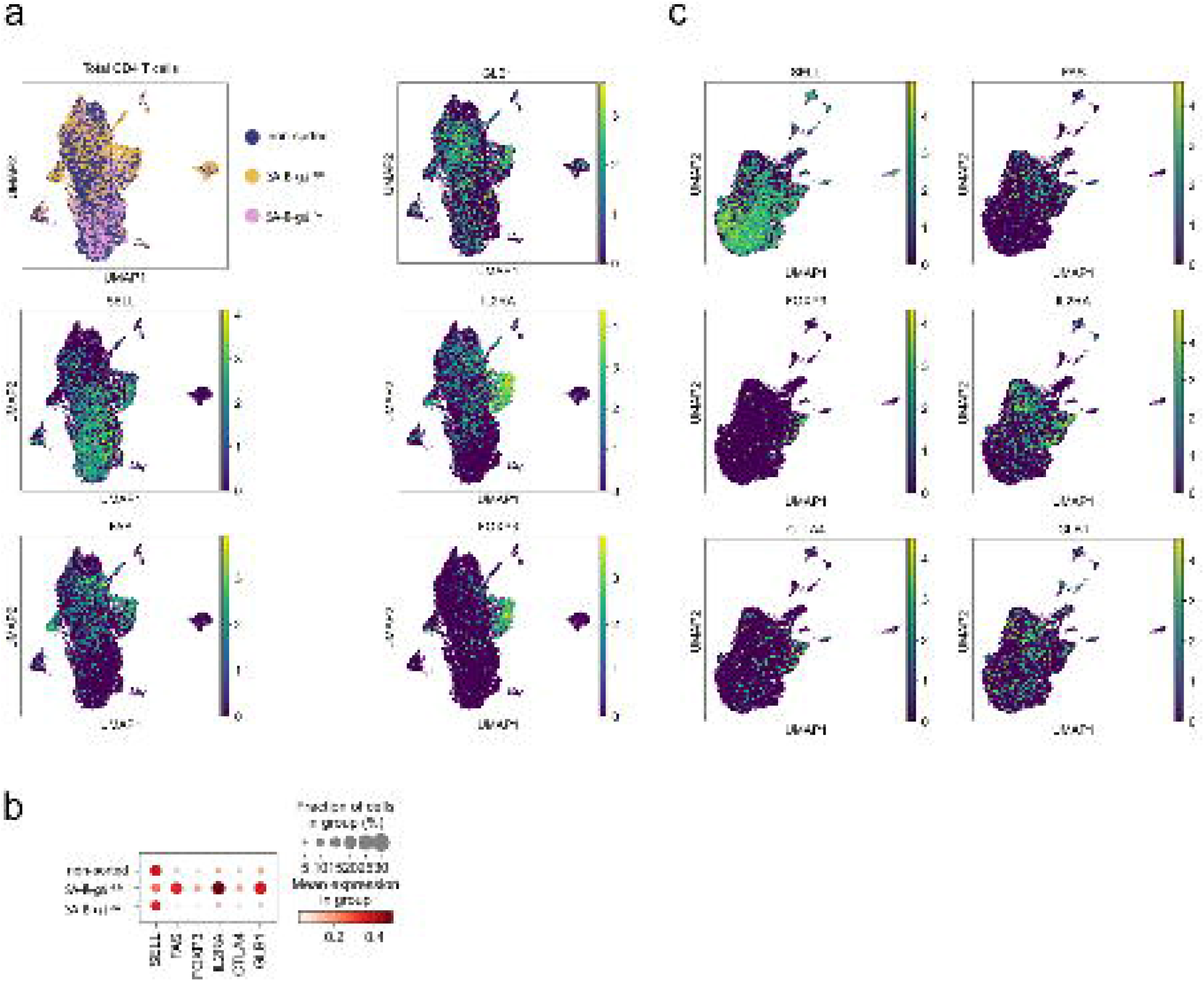
Additional repetitions confirming data reproducibility. **a**,**b**. Similar to Figure 1b,c. Expression of CD4 cell type markers in C_12_FDG-sorted cells. Repeat with independent donor. **c**. Supporting UMAP plots for Figures 1e,f. Naïve CD4 T cells from 15 donors were analyzed without C_12_FDG-sorting. Panels show expression of markers of various CD4 cell types.

**Supplementary Figure S2.**
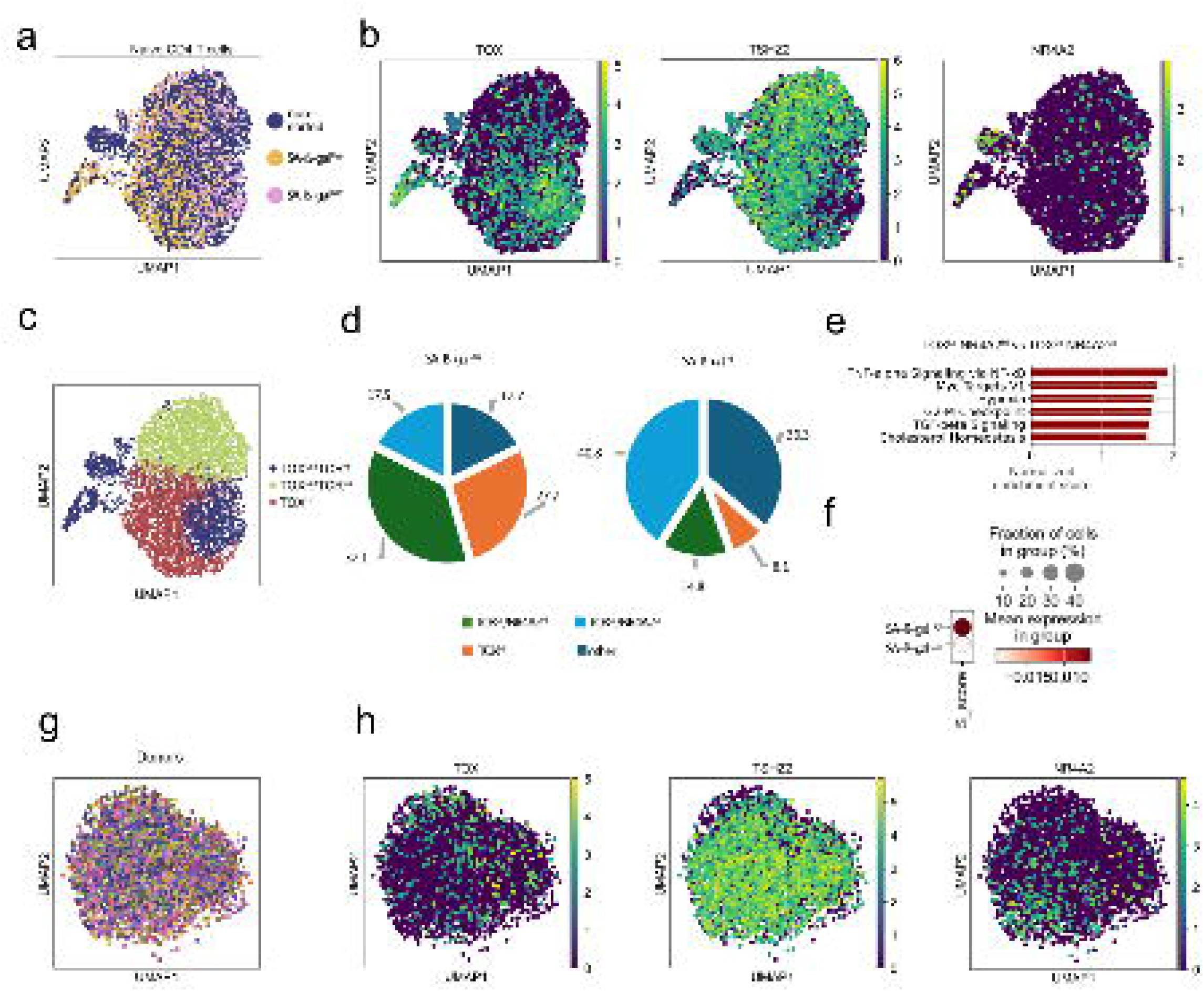
Additional repetitions confirming data reproducibility. **a**. Similar to Figure 2a. UMAP embedding of naïve CD4+ T cells sorted based on C_12_FDG staining. Separate donor from Figure 2a. **b**. Similar to Figure 2c. Expression of the identified marker genes in sorted fractions. **c**. Annotation of the identified subgroups of naïve CD4 T cells. **d**. Abundance of the identified cell groups in SA-β-gal^Hi^ vs. SA-β-gal^Lo^ fractions. **e**. Similar to the first donor, GSEA indicated elevated NF-κB signaling in TOX^Lo^/NR4A2^Hi^ compared to TOX^Lo^/NR4A2^Lo^ cells. **f**. Cell cycle analysis of naïve CD4 T cells 24 hrs after beginning of activation. SA-β-gal^Hi^ showed faster initiation of S phase compared to SA-β-gal^Lo^ cells, consistent with cell division assay shown in Figure 2h. **g**. Naïve cells only from the analysis of multiple donors shown in Figure 1e. **h**. Non-sorted naïve cells from multiple donors show similar heterogeneity characterized by markers TOX, TSHZ2, NR4A2.

